# Homeostasis, injury and recovery dynamics at multiple scales in a self-organizing intestinal crypt

**DOI:** 10.1101/2022.12.18.520934

**Authors:** Louis Gall, Carrie Duckworth, Ferran Jardi, Lieve Lammens, Aimée Parker, Ambra Bianco, Holly Kimko, D. Mark Pritchard, Carmen Pin

## Abstract

We have built a multi-scale agent-based model (ABM) that reproduces the self-organizing behaviour reported for the intestinal crypt. We demonstrate that a stable spatial organization emerges from the dynamic interaction of multiple signalling pathways, such as Wnt, Notch, BMP, RNF43/ZNRF3 and YAP-Hippo pathways, which regulate proliferation and differentiation, respond to environmental mechanical cues, form feedback mechanisms and modulate the dynamics of the cell cycle protein network.

The model recapitulates the crypt phenotype reported after persistent stem cell ablation and after the inhibition of the CDK1 cycle protein. Moreover, we simulated 5-fluorouracil (5-FU)-induced toxicity at multiple scales starting from DNA and RNA damage, which disturbs the cell cycle, cell signalling, proliferation, differentiation and migration and leads to loss of barrier integrity. During recovery, our in-silico crypt regenerates its structure in a self-organizing, dynamic fashion driven by dedifferentiation and enhanced by negative feedback loops.

Overall, we present a systems model able to simulate the disruption of molecular events and its impact across multiple levels of epithelial organization and demonstrate its application to epithelial research and drug discovery.

## Introduction

The intestinal tract is lined by a cellular monolayer which is folded to form invaginations, called crypts, and protrusions, called villi, in the small intestine. The stem cell niche is formed by intermingling Paneth and stem cells located at the base of the crypt (1). Stem cells divide symmetrically, forming a pool of equipotent cells that replace each other following neutral drift dynamics (2). Continuously dividing stem cells at the base of the crypt give rise to secretory and proliferative absorptive progenitors that migrate towards the villus, driven by proliferation-derived forces (3). The transit amplifying region above the stem cell niche fuels the rapid renewal of the epithelium (Figure 1). The equilibrium of this dynamic system is maintained by cell shedding from the villus tip into the gut lumen (4).

**Figure 1.**
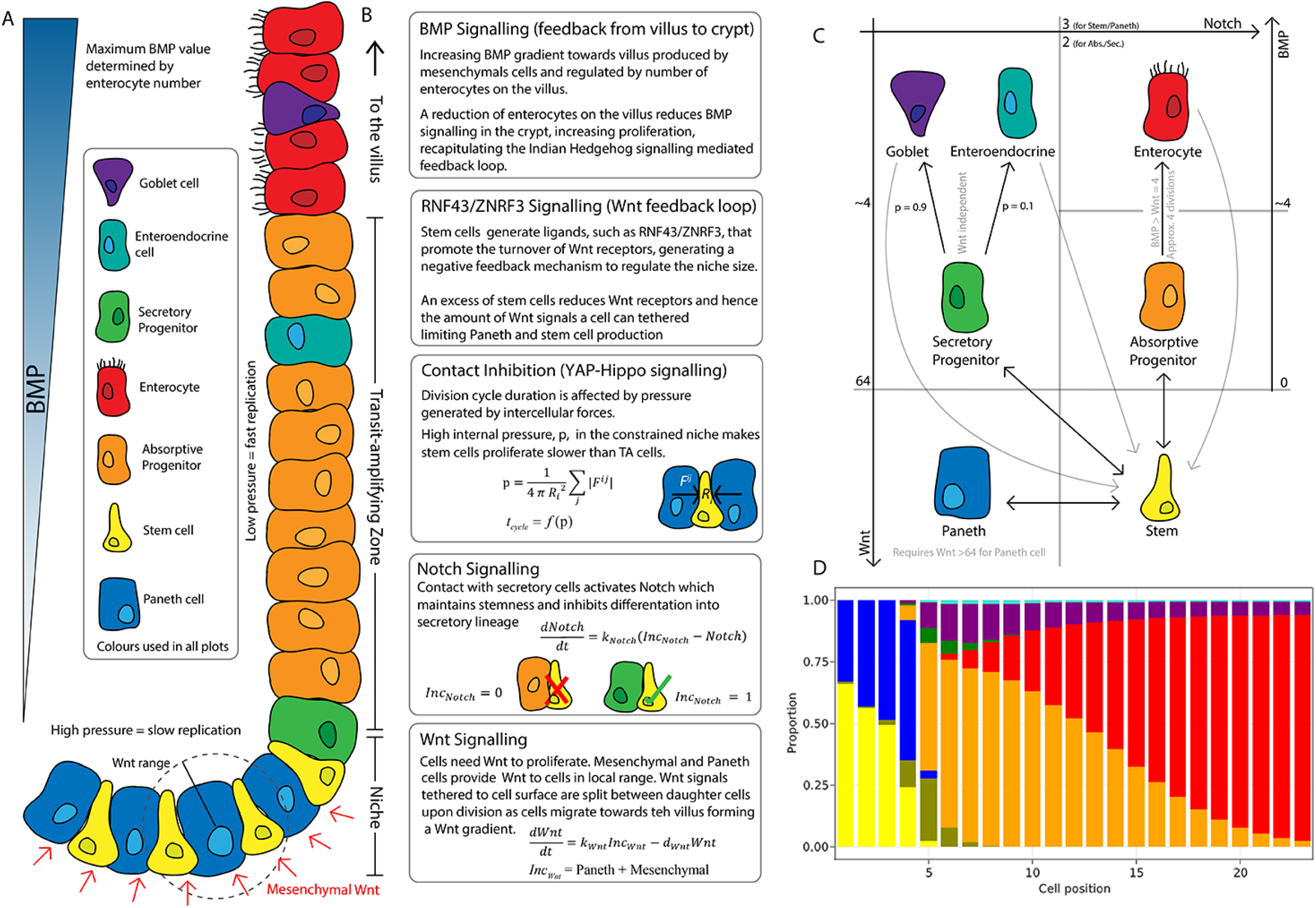
Schematics of the small intestinal crypt composition and cell fate signalling pathways included in the ABM. A) Depiction of the crypt highlighting key signalling features and cell types in each crypt region; B) Details of signalling pathways including high levels of Wnt in the stem cell niche generated by Paneth and mesenchymal cells. Intercellular pressure (contact inhibition of proliferation) regulates the duration of the division cycle which is associated with the accumulation of Wnt signals while Notch signalling maintains the balance between Paneth and stem cells through lateral inhibition. A RNF43/ZNRF3-medited feedback mechanism regulates Wnt signalling in the niche restricting the number of stem and Paneth cells. BMP signals generated by mature villus cells form a feedback loop that regulates maturation and proliferation of absorptive progenitors; C) Cell fate determination. High Wnt signalling and activation of Notch are required to maintain stemness. Low Notch signalling determines differentiation into secretory fates, including Paneth cells in high Wnt signalling regions, or goblet/enteroendocrine progenitors in low Wnt regions. Absorptive progenitors develop from stem cells in low Wnt conditions and divide 3-5 times, depending on BMP signalling, before becoming terminally differentiated; D) Average composition of a simulated healthy/homeostatic crypt (over 100 simulated days), showing the relative proportion of cells at each position in the crypt.

Epithelial cell dynamics is orchestrated by tightly regulated signalling pathways. Two counteracting gradients run along the crypt-villus axis: the Wnt gradient, secreted by mesenchymal and Paneth cells at the bottom of the crypt, and the bone morphogenetic protein (BMP) gradient generated in the villus mesenchyme, with BMP inhibitors secreted by myofibroblasts and smooth muscle cells located around the stem cell niche (5). These two signalling pathways are also the target of stabilizing negative feedback loops comprising the turnover of Wnt receptors (6–9) and the modulation of BMP secretion (10, 11). Paneth cells and the niche surrounding mesenchyme also secrete other proliferation-enhancing molecules such as epidermal growth factor (EGF) and transforming growth factor-α (TGFα) (5). In addition, Notch signalling mediated lateral inhibition mechanisms are essential for stem cell maintenance and differentiation into absorptive and secretory progenitors (5). There is also an increasing awareness of the importance of the mechanical regulation of cell proliferation through the Hippo signalling pathway interplaying with several of the key signals, such as EGF, WNT and Notch, although the exact mechanisms are not currently fully understood (5).

Dysregulated cell division is the key hallmark of cancer and cell division proteins such as cyclin dependent kinases (CDKs) are effective targets for cancer treatment (12). In addition, one of the most common means of targeting the cell cycle is to exploit the effect of DNA-damaging drugs (13). Chemotherapy-induced diarrhea is initiated by the direct or indirect effects of chemotherapeutics on the rapidly dividing epithelial cells in the GI tract (14). With an incidence as high as 80%, it is amongst the primary reasons for dose reductions, delays and cessation of treatment and presents a constant challenge for the development of efficient and tolerable cancer treatments (14). Mechanistic quantitative models can help address the clinical translation of safety data to identify safety concerns and quantify risks in the disease context at early stages of drug development.

Several agent-based models have been proposed to describe the complexity and dynamic nature of the intestinal crypt (15–17). The high performance of computational models and their ability to integrate phenomena at multiple scales have been demonstrated in spatiotemporal simulations of colon cancer initiation and treatment (18) and of intestinal tumorigenesis triggered by Wnt-activating mutations (19). Here, we present a lattice-free agent-based model that describes the spatiotemporal dynamics of single cells in the crypt-villus geometry and implements the main proteins governing the cell cycle as well as mechanisms of DNA damage and repair within each cell. These individual cells interact through multiple signalling pathways to generate a self-organising system that recreates cell composition, compartmentalisation and behaviours observed in the intestinal crypt and villus. We show here that our computational model enables the simulation of perturbations in the niche as well as of drug-induced injury and recovery, driven by clinically relevant exposure to cytotoxic drugs. In our ABM, drug-induced molecular perturbations trigger a cascade of disruptive events spanning across the cell cycle to single cell arrest and/or apoptosis, altered cell migration and turnover and ultimately loss of epithelial integrity.

## Results

### Modelling a self-organizing crypt using an agent based model

We have modelled the intestinal crypt as a self-organizing system where cell dynamics and crypt composition arise from local interactions between single cells and the mesenchyme through signalling pathways without externally imposed behaviours.

The model describes the spatiotemporal dynamics of stem cells and progenitors undergoing division cycles and responding to intercellular signalling to differentiate into Paneth, goblet and enteroendocrine cells and enterocytes (Figure 1A). All cells interact physically and biochemically in the geometry of the crypt. Stem cells intermingle with Paneth cells at the bottom of the crypt and randomly replace each other. Progenitors and mature cells migrate towards the villus driven by proliferation forces (Figure 1A). To achieve a stable crypt cell composition under constant cell renewal dynamics, we have implemented several signalling mechanisms which include the Wnt, Notch and BMP pathways essential for morphogenesis and homeostasis of the intestinal crypt (5, 20–23), the YAP-Hippo signalling pathway responding to mechanical forces and modulating contact inhibition of proliferation (24) and a RNF43/ZNRF3-like mediated feedback mechanism between Paneth and stem cells to regulate the size of the stem cell niche according to experimental reports (6, 7, 25) (Figure 1B).

The Wnt pathway is the primary pathway associated with stem cell maintenance and cell proliferation in the crypt (20, 26). Our model implements two sources of Wnt signals described in the crypt: Paneth cells (27) and mesenchymal cells surrounding the stem cell niche at the crypt base (28). Wnt signalling is modelled as a short-range field around Wnt-emitting Paneth and mesenchymal cells with Wnt signals tethered to receptive cells as previously reported (25, 29). Surface tethered signals are split between daughter cells upon cell division (5, 25), which results in a gradual depletion of tethered Wnt signals as cells divide and migrate towards the villus away from Wnt sources (Figures 1A-1B). Notch signalling is also implemented in the model with Notch ligands expressed by secretory cells binding to Notch receptors on neighbouring cells and preventing them from differentiating into secretory fates in a process known as lateral inhibition that leads to a checkerboard/on-off pattern of Paneth and stem cells in the niche (21). Specifically, in our model, high Wnt and Notch signalling environments are required to maintain stemness. With low Notch and high Wnt signalling, stem cells differentiate into Paneth cells. On the other hand, Notch signalling also mediates the process of Paneth cell de-differentiation into stem cells to regenerate the niche as previously reported (30, 31). Stem cells with decreased levels of Wnt signalling, usually located outside the niche, differentiate into absorptive proliferating progenitors or alternatively into secretory progenitors in the absence of Notch signals (Figure 1C).

In our model, the mechanical environment, captured through the YAP-Hippo signalling pathway (24, 32–34), indirectly interacts with the Notch and Wnt signalling pathways. We recapitulate YAP-mediated contact inhibition of proliferation by using cell compression to modulate the duration of the division cycle which increases when cells are densely squeezed, such as in the stem cell niche, and decreases if cell density falls, for instance in the transit amplifying compartment or in cases of crypt damage (Figures 1A-B). Paneth cells are the main driver behind the differential mechanical environment in the niche, where cells with longer cycles accumulate more Wnt and Notch signals. In agreement with experimental reports (35), Paneth cells are assumed to be stiffer and larger than other epithelial cells, requiring higher forces to be displaced and generating high intercellular pressure in the region. These premises imply that Paneth cells enhance their own production by generating Wnt signals and inducing prolonged division times, which increases stem and Paneth cell production and could lead to unlimited expansion of the niche. To generate a niche of stable size, we implemented a negative Wnt feedback loop that resembles the stem cell production of RNF43/ZNRF3 (6–9) ligands to increase the turnover of Wnt receptors in nearby cells. In our model, when the number of stem cells exceeds the homeostatic value, the increased turnover rate of Wnt receptors impedes Paneth and stem cell generation by reducing the capability of cells to acquire sufficient Wnt ligands. This Wnt-mediated feedback loop prevents the uncontrolled expansion of the niche (Figures 1A-B).

The Wnt gradient in the crypt is opposed by a gradient of bone morphogenic protein (BMP) that inhibits cell proliferation and promotes differentiation (36). We assume that enterocytes secrete diffusing signals, resembling Indian Hedgehog signals (10), that induce mesenchymal cells to generate a BMP signalling gradient effective to prevent proliferative cells from reaching the villus (Figures 1A-B). Based on experimental evidence, we also assume that BMP activity is counteracted by BMP antagonist-secreting mesenchymal cells surrounding the stem cell niche (37). Proliferative absorptive progenitors migrating towards the villus lose Wnt during every division and eventually meet values of BMP that overcome the proliferation-inducing effect of Wnt signalling (23). We found that an optimal crypt cell composition is achieved when BMP and Wnt differentiation thresholds result in progenitors dividing approximately four times before differentiating into enterocytes (Figure 1C). In addition, the BMP signalling gradient responds dynamically to the number of enterocytes, giving rise to a negative feedback loop between enterocytes on the villus and their proliferative progenitors in the crypt that recapitulates the enhanced crypt proliferation observed after epithelial damage (10, 38, 39). For instance, a decreased number of enterocytes results in reduced production of BMP, which enables progenitor cells to divide and migrate further up the crypt before meeting BMP levels higher than the differentiation threshold.

All together our model describes single cells that generate and respond to signals and mechanical pressures in the crypt-villus geometry to give rise to a self-organizing crypt which has stable cell composition over time (Figure 1D). An extended description of these modelling features is provided in the *Apendix*.

### The cell cycle protein network governs proliferation in each single cell of the ABM and responds to the mechanical environment

We have used the model of Csikasz-Nagy *et al*. (40), available in BioModels (41), to recreate the dynamics of the concentration of the main proteins governing the mammalian cell cycle in each single proliferative cell of the ABM. In this model, a dividing cell begins in G1, with low levels of Cyclins A, B and E and a high level of Wee1, and progresses to S-phase when Cyclin E increases. S-phase ends and G2 begins when Wee1 falls. The decrease in Cyclin A expression defines the start of M-phase, while falling Cyclin B implies the end of M-phase, when the cell divides into two daughter cells with half the final mass value and re-enters the cell cycle (Figures 2A-2D).

**Figure 2.**
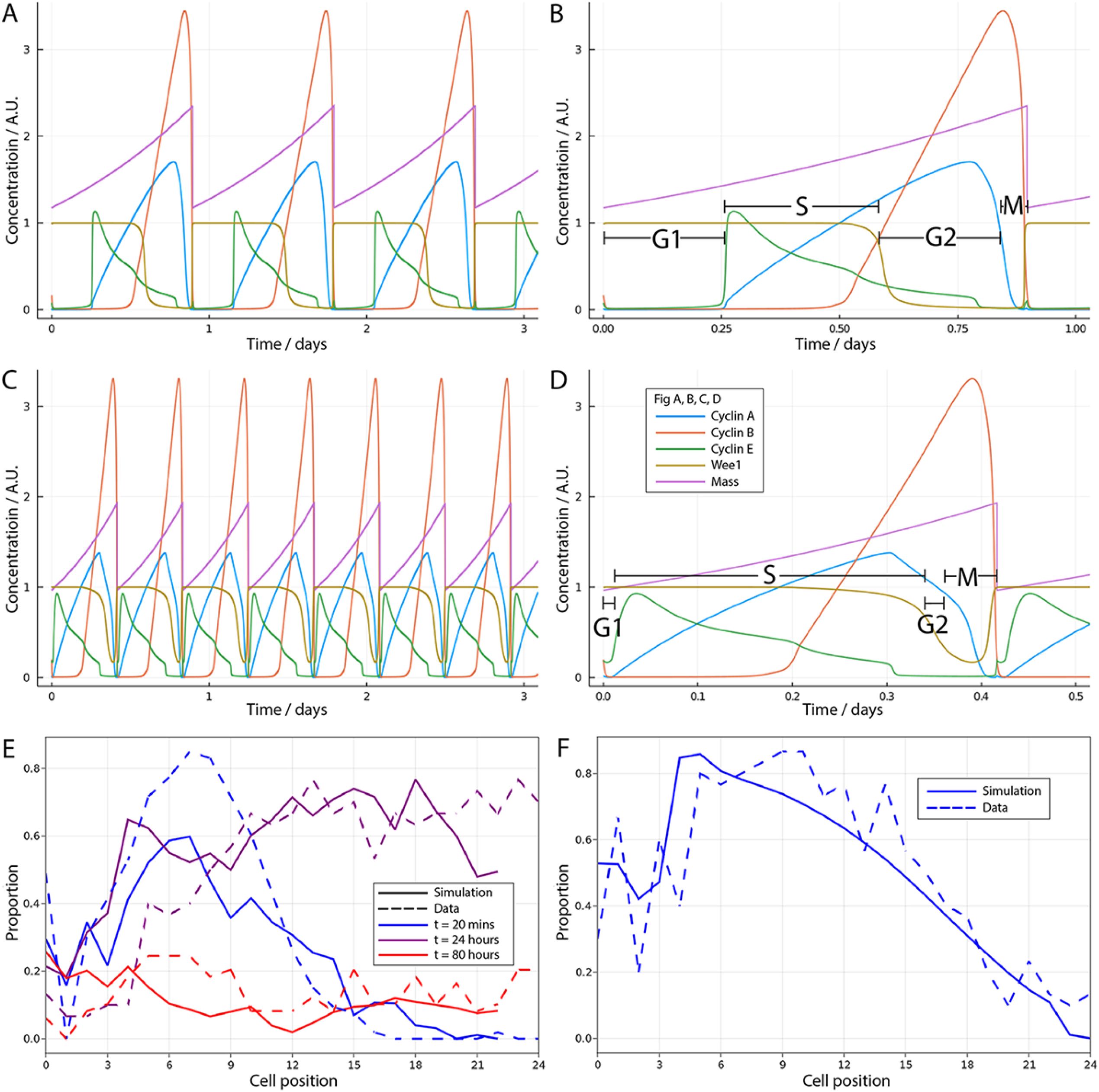
Modelling cell division in single cells of the ABM. A-D) Modelled oscillatory proteins (40) across cell cycle stages (B,D) and during repeated division cycles (A,C) of stem cells (A,B) and faster cycling transit amplifying cells (C,D) with shortened G1 phase, with concentrations given in arbitrary units (A.U.); E) Observed (dashed line) and simulated (solid line) proportions of BrdU positive cells at each crypt position at 20 mins (blue), 24 hours (purple) and 80 hours (red) after a single pulse of BrdU; F) Observed and simulated Ki-67 positive cells at each crypt position assuming that Ki-67 is detected in cycling cells at all phases except G1 and in any recently differentiated and arrested cells that were cycling within the last 6 hours (47).

To implement YAP-Hippo mediated contact inhibition of proliferation, we have modified the dynamics of the proteins of the Csikasz-Nagy model to respond to the mechanical environment encountered by cells migrating along the crypt. Crowded, constrained environments result in longer cycles, such as in stem cells in the niche, while decreased intercellular forces lead to shortened cycles as cells migrate towards the villus in agreement with experimental reports (4, 42, 43). The shorter cycle duration in absorptive progenitors has been mainly associated with shortening/omission of G1, while the duration of S phase is less variable (4). Using the model of Csikasz-Nagy *et al*. (40), we modulated the duration of G1 through the production rate of the p27 protein, which prevents the activation of Cyclin E-Cdk2 required to start DNA replication and S-phase (44). In the modified model, low levels of p27 bring forward the start of S-phase and shorten G1 in cells experiencing low intercellular pressure (Figures 2D). In support of this hypothesis, it has been demonstrated that p27 inhibition had no effect on the proliferation of absorptive progenitors (45). In addition, to regulate the duration of S-phase, we used the parameter governing the phosphorylation of Wee1, which controls the onset of G2-phase and the end of S-phase (see the *Appendix* for a full description). These new features of the cell cycle model are updated dynamically and continuously to respond to changes in mechanical pressure experienced by each cell as it migrates along the crypt.

To demonstrate the performance of the model in homeostatic conditions, we simulated previous published mouse experiments (3, 46) comprising BrdU tracking (Figure 2E) and Ki-67 staining (Figure 2F). To simulate the BrdU chase experiment after a single BrdU pulse, we assumed that any cell in S-phase, incorporated BrdU permanently into its DNA for the first 20 minutes after dosing and BrdU cell content, was diluted upon cell division such that after five cell divisions, BrdU was not detectable. The BrdU chase experiment showed that the observed initial distribution of cells in S-phase as well as division, differentiation and migration of BrdU-positive cells over time were replicated by our model (Figure 2E). Regarding the Ki-67 proliferative marker, our simulations assumed that Ki-67 was detected in cycling cells at all phases except G1 as well as in recently arrested or differentiated cells that were cycling within the past 6 hours (47). Similarly, we observed that the ABM-simulated spatial distribution along the crypt of Ki-67 positive cells recapitulated observations in mouse jejunum (Figure 2F).

In summary, the proliferative cells in the ABM respond to mechanical signals by adjusting the cell cycle protein network to dynamically change the duration of the cycle while migrating along the crypt. With this feature, the model replicates the spatiotemporal patterns of cell proliferation, differentiation and migration observed in mouse experiments.

### Cell plasticity/de-differentiation enables crypt regeneration following damage of the stem cell niche

Marker-based lineage tracing studies have demonstrated numerous potential sources available for intestinal stem cell regeneration (48). In line with these studies, our model assumes that cell fate decisions are reversible and both secretory and absorptive cells are able to revert into stem cells when regaining sufficient Wnt and Notch signals.

To investigate the potential of the ABM to describe and explore cell plasticity dynamics, we simulated the repeated ablation of intestinal stem cells resembling a previously published study (49). Following the experimental set up in that study, we simulated the diphtheria toxin receptor-mediated conditional targeted ablation of stem cells for four consecutive days by persistently inducing stem cell death in a process completed after 24h (50) (Figure 3A-C). Our simulations showed that 6 hours after the last induction, stem cells were not detected, Paneth cells decreased by 75% (Figure 3B) and the villus length was reduced by about 10-20% (Figure 3C) which was similar to the reported experimental findings (49). Simulated proliferative absorptive progenitors initially decreased, to reach minimum values by day three, but their recovery started before the end of the treatment, driven by a negative feedback loop from mature enterocytes to their progenitors, to later reach values above baseline (Figure 3A). Hence, in our simulations, enhanced crypt proliferation was not accompanied by simultaneous villus recovery, which started later. Tan *et al*. (49) reported similar results with increased crypt proliferation replenishing first the crypt and not contributing immediately to villus recovery. See *Supplementary Movie 1* to visualize the response of the crypt.

**Figure 3.**
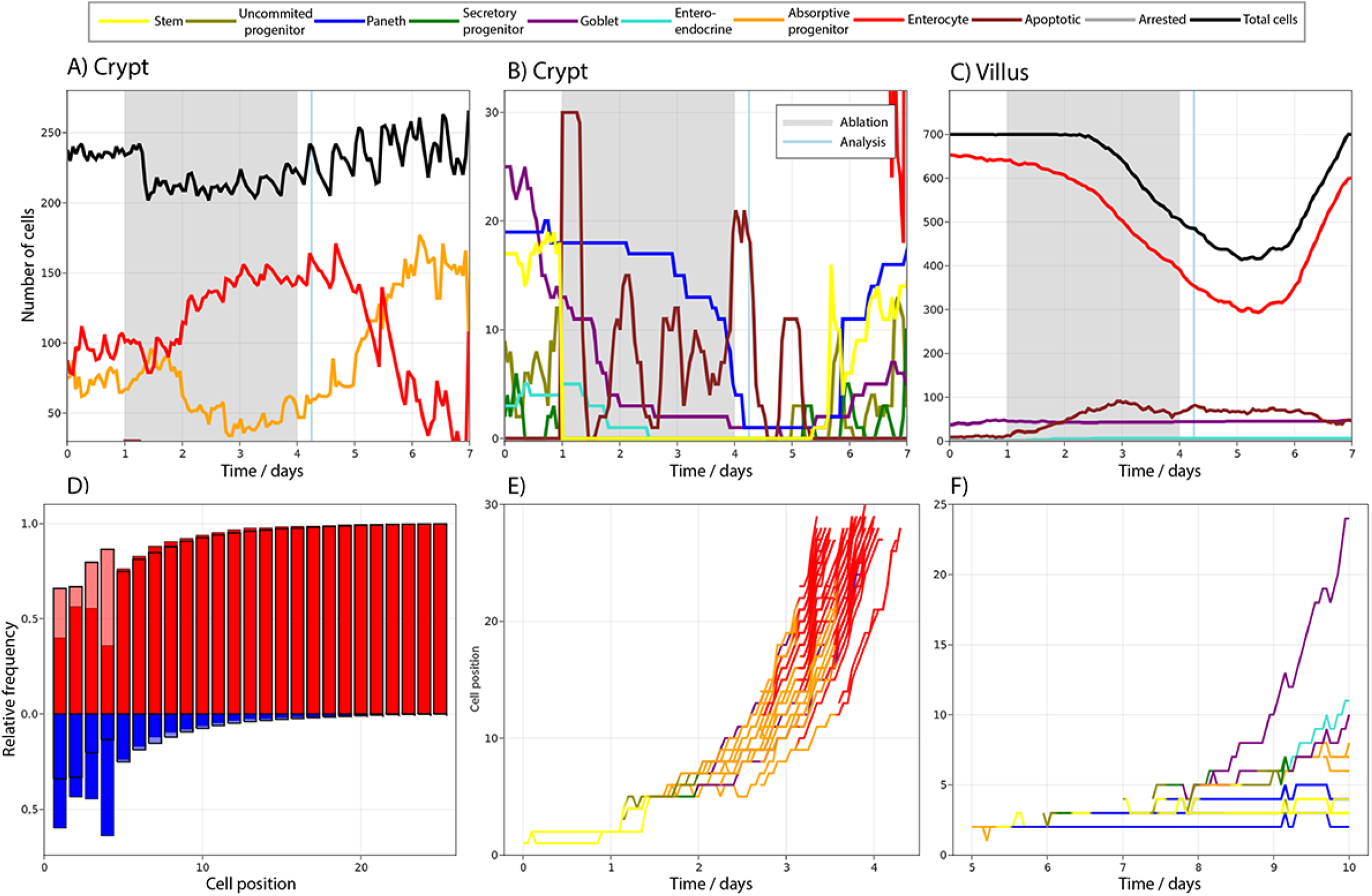
A-C) Simulated cell dynamics in the epithelium subjected to continuous ablation of stem cells for 4 consecutive days (grey block) resembling a previously published experiment (49). All cell lineages are recorded during treatment and recovery in the simulated crypt: (A) shows the most abundant cell types in the crypt, (B) shows crypt cells with lower counts and (C) shows villus cells. D) relative frequency of crypt cells moving upwards, towards the villus (red), or downwards, towards the crypt base (blue), in homeostasis (black outlined columns) at each cell position and during stem cell ablation (no outlined columns), showing increased retrograde cellular motion as cells repopulate the niche. E) Trajectory of one stem cell progeny in homeostasis, with both daughters leaving the niche and proliferating into a cascade of absorptive and secretory cells that eventually leave the crypt. F) Trajectory of the progeny of an absorptive progenitor returning to the niche following the cessation of the stem cell ablation. The cell dedifferentiates into a stem cell that regenerates multiple stem and Paneth cells in the niche, as well as secretory and absorptive progenitors that move upwards.

We next studied the type of cells that were dedifferentiating during the simulated repeated ablation of stem cells and found that in agreement with experimental reports, Paneth cells (31), absorptive progenitors (51) and + 4 quiescent stem cells (52) dedifferentiated into stem cells. Specifically, from all dedifferentiated cells, about 60% were Paneth cells, 30% absorptive progenitors and 10% secretory progenitors, which could be considered quiescent +4 stem cells as previously suggested (53). Furthermore, we used our model to explore the retrograde motion, reported using intravital microscopy (54), of cells returning to the niche to de-differentiate into stem cells. For cells outside the niche, movement is retrograde when its velocity is negative in the *z* direction. For cells in the hemispherical niche, we consider a cell to move forward, towards the villus, or backward, towards the crypt base, if the rate of change of its polar angle is positive or negative, respectively. This implies that cells can be recorded to move backwards despite being located at the crypt bottom. We observed that the frequency of retrograde, or backward, movements is relatively high at low positions in a crypt in homeostasis (Figure 3D) and increases further after stem cell ablation, reflecting increased retrograde cellular motion as cells repopulate the niche. While in homeostasis the progeny of a stem cell generally differentiate into a cascade of absorptive and secretory progenitors that eventually leave the crypt (Figure 3D), following the interruption of stem cell ablation, absorptive progenitors return to the niche and dedifferentiate to regenerate multiple stem and Paneth cells as well as progenitors (Figure 3F).

Taken together, our model recapitulates cellular reprogramming of both multipotent precursors and committed progeny in the crypt and replicates the reported crypt injury dynamics following persistent ablation of stem cells (49).

### Disturbance of cell cycle proteins spans across scales to impact on crypt and villus organization

The model of Csikasz-Nagy *et al*. (40) enables the simulation of the disruption of the main proteins governing the cell cycle in each single proliferative cell of the ABM. CDKs play important roles in the control of cell division (55) and the development of CDK inhibitors for cancer treatment is an active field of research (12).

To explore the effect of the disruption of the cell cycle on epithelial integrity, we simulated the inhibition of CDK1, which is reported to be the only CDK essential for the cell cycle in mammals (56). CDK1 triggers the initiation of cytokinesis by inducing the nuclear localization of mitotic cyclins A and B (57) and its inhibition has been proposed as a cancer therapy with potentially higher efficacy than the inactivation of other CDKs (58). To mimic CDK1 inhibition, we added a term to the CycA/CDK1,2 and CycB/CDK1 differential equations of the Csikasz-Nagy model (40) that strongly reduces the production of both CycA/CDK1,2 and CycB/CDK1 for 4 consecutive days, resembling the effect of the administration of a theoretical drug (Figure 4 A-E) (See the *Appendix*).

**Figure 4.**
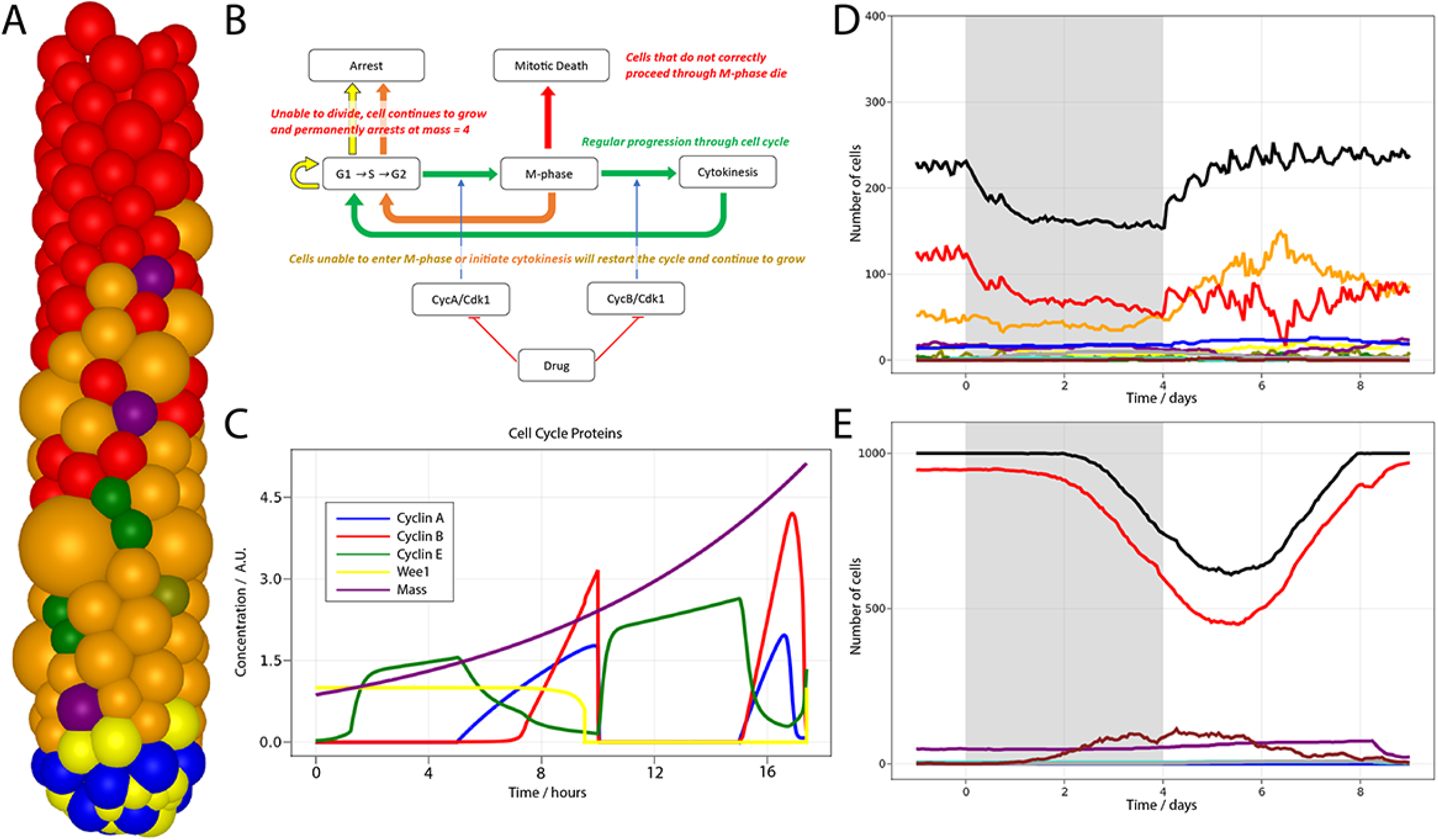
Simulation of CDK1 inhibition in the ABM and impact on the cell cycle and crypt and villus organization. A) Following CDK1 inhibition, the crypt ABM exhibits over-sized cells that are unable to correctly complete the cell cycle; B) Flowchart showing the regular progression through the cell cycle (green path) disturbed by CDK1 inactivation. Cells can fail to culminate mitosis, but restart G1 and the cycle when CDK1 inactivation prevents either cells at the end of G2 from entering M-phase (yellow path) or cells in M-phase from completing cytokinesis (orange path), or spend insufficient time in M-phase and undergo mitotic death (red path); C) Altered cell cycle protein profile disturbing G2 and M-phase and preventing the cell mass from dividing before reinitiating a new cycle, with concentrations given in arbitrary units (A.U.); Cell dynamics in simulated crypts (D) and villi (E) is affected by CDK1 inhibition. Colour scheme in D and E as in Figures 1 and 3.

It has been experimentally demonstrated that the selective inhibition of CDK1 activity in cells programmed to endoreduplicate (i.e. cells that can duplicate their genome in the absence of intervening mitosis) leads to the formation of stable nonproliferating giant cells, whereas the same treatment triggers apoptosis in cells that are not developmentally programmed to endoreduplicate (59). Although endoreduplication is not expected in crypt cells, enlarged polynucleated cells have been reported to remain in the epithelium without dying in a recent light-sheet organoid imaging study tracking the progeny of a cell after cytokinesis failure, which was induced by the inhibition of LATS1 (60), which is a phosphorylated by CDK1 during mitosis (61). Thus, we chose to replicate this phenotype to show the capacity of our model to predict possible complex responses in the intestine. Following CDK1 inhibition, we detected over-sized cells in the ABM (Figure 4A). The inhibition of the activation of cyclins A and B did not impact on S-phase but altered the modelled protein profiles disturbing G2 and M-phase and preventing the cell mass from dividing before reinitiating a new cycle (Figure 4B). Thus, a cell could either be (i) unaffected if it was at the early stages of the cycle, or (ii) restart G1 and the cell cycle if CDK1 was inhibited while the cell was at the end of G2 and unable to enter M-phase or (iii) in M-phase and unable to complete cytokinesis (Figure 4C). The failure to culminate M-phase resulted in the generation of over-sized, nonproliferating cells, which led to a reduction of the crypt overall cell number (Figure 4D) and the turnover of villus cells (Figure 4E). See *Supplementary Movie 2* to visualize the response of the crypt to CDK1 inhibition.

Altogether our ABM enables the simulation of how disruptions of the cell cycle protein network span across scales to generate complex phenotypes, such as giant cells, and impact on the integrity of the crypt and villus structure.

### A practical application of the ABM to describe 5-fluoruracil (5-FU) induced epithelial injury at multiple scales

5-fluoruracil (5-FU) is a well-studied and commonly administered cancer drug (62) with reported high incidence of gastrointestinal adverse effects in treated patients (63). 5-FU is a pyrimidine antimetabolite cytotoxin which has multiple mechanisms of action upon conversion to several nucleotides that induce DNA and RNA damage (62). Antimetabolites resemble nucleotides and nucleotide precursors that inhibit nucleotide metabolism pathways, and hence DNA synthesis, as well as impair the replication fork progression after being incorporated into the DNA (13).

To explore the performance of our ABM to predict epithelial injury, we used partially published data from experiments in mice dosed with 50 and 20 mg/kg of 5-FU every 12 h for four days to achieve drug exposures similar to those observed in patients (64). 5-FU pharmacokinetics (PK) was modelled considering that 5-FU is metabolized into three active metabolites FUTP, FdUMP and FdUTP (62) as described in a previous published report (65). The 5-FU PK model (64) was integrated into the ABM to describe the dynamic profile of the concentration of 5-FU and its metabolites in plasma and GI epithelium after dosing (Figure 5A). Based on previous reports, we assumed that FUTP is incorporated into RNA of proliferative cells leading to global changes in cell cycle proteins and may result in p53-mediated apoptosis(66) while FdUTP is incorporated into DNA (62) during S-phase resulting in the accumulation of damaged DNA which could be repaired or lead to cell arrest (Figure 5B). We did not include the inhibition of thymidylate synthase (TS) by FdUMP because the impact of this mechanism on intestinal toxicity is not completely clear (66).

**Figure 5.**
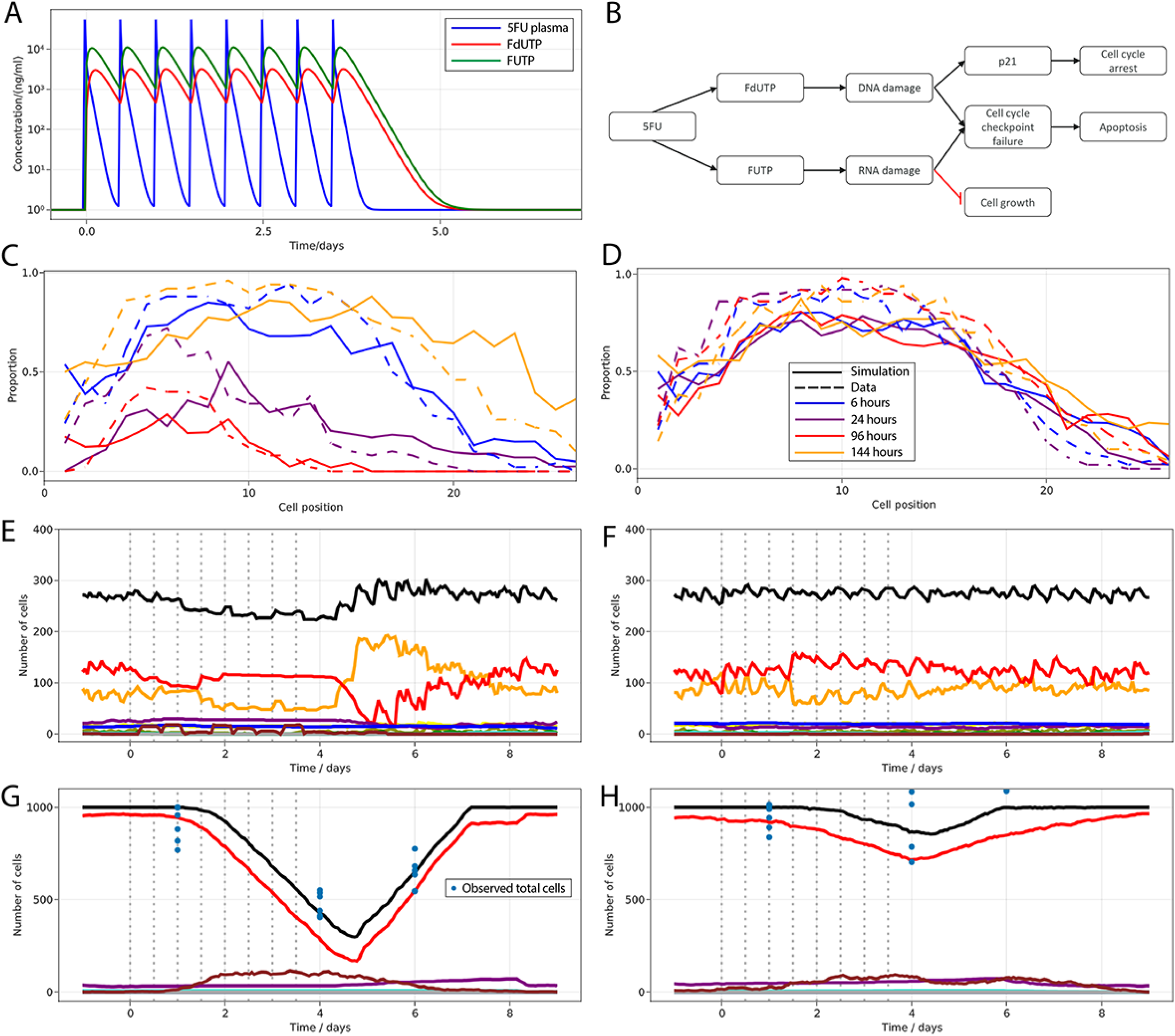
Modelling 5-FU induced injury at several scales in mouse small intestinal epithelium. A) Predicted concentration (ng/ml) of 5-FU in plasma and of 5-FU, FUTP and FdUTP in epithelial cells following the administration of 20 and 50 mg/kg of 5-FU twice a day for four days in mouse (Pharmacokinetics model of 5-FU described in Gall *et al*. (65)); B) Diagram showing the implemented mechanism in the ABM to describe DNA and RNA damage and cell cycle disruption driven by 5-FU metabolites; (C, D) Predicted (solid line) and observed (dashed line) proportions of proliferative cells along the crypt axis at 6h, 1d, 4d and 6d; (E,F) Predicted (lines) number of cells in crypt; (G,H) Predicted (lines) and observed (symbols) number of cells in villus following the administration of 50mg/kg (C, E, G) and 20 mg/kg (D, F, H) of 5-FU twice a day (dosing time indicated by vertical bars) for 4 days. Colour scheme in E, F, G and D as in Figures 1 and 3. The blue circles in G and H are cell counts from individual mice.

Figure 5C shows that predicted and observed Ki-67 positive cells declined gradually over time at all positions in the crypt during the treatment but reached full recovery two days after the interruption of 5-FU administration. The predicted total number of cells in the crypt followed the same pattern in agreement with the recorded data, while the decline of villus cells started later and did not achieve full recovery by the end of the experiment (Figure 5D). Observations showed that the proliferative crypt compartment exhibited a feedback-mediated rebound during recovery, reaching values above baseline, or initial value, (Figure 5C-D) which was captured in our ABM by the BMP-mediated feedback from mature enterocytes to proliferative cells See *Supplementary Movies 3 and 4* to visualize cell dynamics and changes in the main crypt signalling pathways during 5-FU challenge.

Overall, the ABM recapitulates DNA and RNA damage resulting in cell cycle disruption associated with 5-FU administration and describes the propagation of the injury across scales to disturb epithelial integrity. The loss of epithelial barrier integrity is widely accepted to be the triggering event of chemotherapy-induced diarrhea (14) which is reported in mice at the doses used in this study (64) as well as observed in patients undergoing equivalent treatments (67).

## Discussion

We have built a multi-scale agent-based model of the small intestinal crypt with self-organizing stable behaviour that emerges from the dynamic interaction of Wnt, Notch, BMP and RNF43/ZNRF3 pathways orchestrating cellular fate, contact inhibition modulating proliferation and the cell cycle protein interaction network regulating progression across division phases.

In our model, the stability of the niche is achieved by a negative feedback mechanism from stem cells to Wnt respondent cells that resembles the reported turnover of Wnt receptors by RNF43/ZNRF3 ligands secreted by stem cells (6–9). Wnt signals generated from mesenchymal cells and Paneth cells at the bottom of the crypt form a decreasing gradient towards the villi that stimulates cell proliferation and stemness maintenance (25, 27). The BMP signalling counter-gradient along the crypt-villus axis is simulated, resembling BMP production by mesenchymal telocytes abundant at the villus base as well as BMP antagonist secretion by trophocytes located just below crypts (37). An additional negative-feedback mechanism implemented in our model regulates the size of the crypt proliferative compartment and recapitulates the modulation of BMP secretion by mesenchymal cells via villus cells-derived hedgehog signalling (10, 11).

Another novel feature of our model is the inclusion of the dynamics of the protein network governing the phases of cell division (40). Moreover, our cell cycle protein network responds to the mechanical environment by adapting the duration of the cycle phases. Cells in crowded environments subjected to higher mechanical pressure, such as stem cells in the niche, exhibit longer cell cycles (4, 42, 43) while progenitors in the transit amplifying compartment adapt their cell cycle protein dynamics to mainly shorten G1 phase (4, 68) and proliferate more rapidly. This model feature recapitulates the widely reported YAP-mediated mechanism of contact inhibition of proliferation under physical compression (32–34). Interestingly, it has been reported that stiff matrices initially enhance YAP activity and proliferation of *in-vitro* cultured intestinal stem cells by promoting cellular tension (24), however, that study also proposes that the resulting colony growth within a stiff confining environment may give rise to compression YAP inactivation retarding growth and morphogenesis (24).

Furthermore, our model considers that the mechanical regulation of the cell cycle interacts with signalling pathways to maintain epithelial homeostasis, but also to trigger cell dedifferentiation if required. Cells with longer cycles accumulate more Wnt and Notch signals, leading to the maintenance of the highly dynamic niche by replacement of Paneth and stem cells. Cells located outside the niche exhibit shorter cycles and cannot effectively accumulate enough Wnt signals to dedifferentiate into stem cells in homeostatic conditions. However, in case of niche perturbation, progenitor cells reaching the niche as well as existing Paneth cells in the niche are able to dedifferentiate into stem cells after regaining enough Wnt signals, which replicates the injury recovery mechanisms observed in the crypt (48, 51). Our model also concurs with experimental results suggesting that Lgr5+ stem cells are essential for intestinal homeostasis and that their persistent ablation compromises epithelial integrity (49).

Altogether, our model implements qualitative and quantitative behaviours to better simulate the functional heterogeneity of the intestinal epithelium at multiple scales. One of the important applications of our modelling approach lies in the discovery of safer oncotherapeutics. Here we demonstrated the application of our model to predict potential intestinal toxicity phenotypes induced by CDK1 inhibition as well as to describe the disruption of the epithelium at multiple scales triggered by RNA and DNA damage leading to the loss of integrity of the intestinal barrier and diarrhea following 5-FU treatment. Thus, the opportunity emerges to assess the safety of therapeutic agents early during drug development. Our work highlights the importance of novel modelling solutions that are able to integrate the dynamics of processes at multiple scales regulating the functionality of the intestinal epithelium in homeostasis and following perturbations to provide unprecedented insights into the biology of the epithelium with practical application in the development of safer novel drug candidates.

## Materials and Methods

### Mouse experiments

BrdU tracking and Ki-67 immunostaining data were derived from previously published experiments in healthy mice (3, 46) and following 5-FU treatment (64). Ki-67 immunostaining data during 5-FU treatment was derived from the same published experiments (64). However, the previously reported data included overall cell counts per crypt and villus, whereas here we use the counts of Ki-67 positive cells recorded by position along the longitudinal crypt axis, for 30-50 individual hemi crypt units per tissue section per mouse (69).

### Agent based model development

A comprehensive description of the model can be found in the *Appendix* and *Supplementary Table 1*. The model has been made available through BioModels (MODEL2212120003) (70)

## Acknowledgments

The authors acknowledge financial support from TransQST consortium. This project has received funding from the Innovative Medicines Initiative 2 Joint Undertaking under grant agreement No 116030. This Joint Undertaking receives support from the European Union’s Horizon 2020 research and innovation programme and EFPIA.

## Conflict of interest

LG, AB, HK and CP are employees and shareholders of AstraZeneca Plc. LL and FJ are employees of Johnson & Johnson. LL is a shareholder of Johnson & Johnson.

## Supplementary movie caption

**Supplementary Movie 1.** Simulated cell dynamics in the epithelium subjected to continuous ablation of stem cells for 4 consecutive days resembling a previously published experiment (49). Plots depict changes in the number of cells in the crypt and villus during the simulation. Colour code of cell types described in Figures1-5.

**Supplementary Movie 2.** Simulated cell dynamics in the epithelium subjected to CDK1 inhibition for 4 days. Plots depict changes in the number of cells in the crypt and villus during the simulation. Colour code of cell types described in Figures1-5.

**Supplementary Movie 3.** Simulated cell dynamics in the epithelium following the administration of 50 mg/kg of 5-FU twice a day for four days in mouse (Pharmacokinetics model of 5-FU described in Gall *et al*. (65)). Plots depict changes in the number of cells in the crypt and villus during the simulation. Colour code of cell types described in Figures1-5.

**Supplementary Movie 4.** Simulated main epithelial signalling pathways following the administration of 50 mg/kg of 5-FU twice a day for four days in mouse (Pharmacokinetics model of 5-FU described in Gall *et al*. (65)). Plots depict changes in signal abundance across the crypt longitudinal axis (*z*) during the simulation. Colour code of cell types described in Figures1-5.

## Appendix. Technical description of the intestinal epithelial agent-based model (ABM)

The model primarily focuses on describing the spatiotemporal dynamics of single epithelial cells, interacting physically and biochemically in the intestinal crypt, undergoing division cycles or differentiating into mature epithelial cells. Single cells both generate and respond to signals and mechanical pressure in the crypt-villus geometry to generate a self-organizing tissue.

Below we describe the assumptions and hypotheses that underpin the model, regarding 1) geometry; 2) cell cycle proteins and cellular growth; 3) drug perturbation of the cell cycle proteins: Cdk1 inhibition; 4) DNA and RNA synthesis; 5) drug perturbations of RNA and DNA synthesis: 5-FU induced RNA and DNA damage; 6) mechanical cell interactions and contact inhibition; 7) biochemical signalling; 8) cell fate: proliferation, differentiation, arrest, apoptosis; 9) model implementation and parameterization.

### 1) Geometry

To recreate the morphology of the crypt, we chose the common idealised ‘test tube’ crypt geometry of a hemisphere attached to a cylinder, which describes the basement membrane that the cells are attached to. The parameters describing the average morphology of the crypt, i.e. the height and circumference of the ‘tube’, in mouse are described in Supplementary Table 1.

Cells on the villus are terminally differentiated and can be assumed to migrate on a conveyor belt at constant velocity (3). Given these simple dynamics, to save computational power and time we modelled individual cells on the villus without spatial granularity. Cells that reach the top of the crypt are collected into a villus compartment. Shedding from the villus tip is mimicked by removing the oldest cells when the number of cells exceeds the maximum capacity of the villus, which is described in Table 1. Cells on the villus keep all properties and still age and undergo apoptosis if required, though in homeostatic conditions cells are usually shed into the lumen before becoming senescent.

### 2) Cell cycle proteins and cellular growth

The division cycle of cells is controlled by a network of interacting proteins which include cyclins, cyclin-dependent kinases (CDKs) and a suite of ancillary proteins (44). The discrete events of the cell cycle, such as DNA replication in S-phase and the various stages of mitosis, are regulated by the activity of this protein network, whose components go through a careful, conserved series of peaks and troughs at the correct pace to complete all processes of the cycle. The dynamics of this protein interaction network is simulated in each cell of the ABM and controls cell division and differentiation.

We have used the model of Csikasz-Nagy *et al*. (40), that recreates the mammalian cell cycle and is available in Biomodels (41). The model compromises 14 variables that describe the dynamics of the concentration of the main cell cycle proteins as oscillations between alternating peaks and troughs. G1 phase is the default opening state, with low levels of Cyclins A, B and E and high level of Wee1. The level of cyclin D grows exponentially throughout the cycle and is halved between daughter cells after mitosis. S-phase begins with the increase of Cyclin E and ends when Wee1 drops to reach its trough. G2 phase is characterised by low Wee1 and high Cyclin A, ending with the drop of Cyclin A. M-phase ends when Cyclin B falls and the cell divides and restarts the cycle in G1.

Stem cells have been reported to have a longer division cycle than absorptive progenitor cells (4), (42), (43). We hypothesise that this is due to contact inhibition mechanisms caused by increased intercellular forces in the crowded, constrained niche. This implies that the duration of the cycle may significantly vary among single cells. To implement cycles of varying duration in our ABM we describe below a series of required adjustments in the Csikasz-Nagy model that basically involve changes in the duration of the full cycle, the re-adjustment of the length of the cycle phases, primarily G1 and S phase, and the modulation of the dynamics of the model mass variable.

To change the duration of the cell cycle, *t_cycle_*, we rescaled the time coordinate: 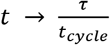 *t*, where *τ* = 140.027 h is the original period of the model (40) and *t_cycle_* is determined by the internal pressure of the cell as detailed below in the section “Mechanical Cell Interactions”.

Without further modifications of the Csikasz-Nagy model (40), the duration of all cycle phases would be scaled in proportion with changes in *t_cycle_*. However, not all phases are proportionally shortened in fast cycling healthy cells (40). The shorter cycle duration in absorptive progenitors is likely due to shortening/omission of G1 phase as reported for rapid cycling progenitors (4, 68), while the duration of S-phase is less variable (4) with reported values of 8 hours for mouse ileal epithelium (4).

Regarding G1 phase, the p27 protein has been reported to regulate the duration of G1 by preventing the activation of Cyclin E-Cdk2 which induces DNA replication and defines the beginning of S-phase (44). We hypothesized that fast cycling cells have low levels of p27 which result in earlier DNA replication, bringing forward the start of S-phase and shortening the length of G1. In support of this hypothesis, it has been experimentally demonstrated that inhibiting p27 has no effect on the proliferation of absorptive progenitors (45). In the Csikasz-Nagy model (40), the duration of G1 can be modulated through the parameter *V_si_*, which is the basal production rate of p21/p27 (in the Csikasz-Nagy model, the p21 and p27 proteins are represented by a single variable, here we refer to that model quantity as p21/p27).

Additionally, the end of S-phase is associated with the decrease of Wee1 to basal levels due to Cdc14 mediated phosphorylation of Wee1. In the Csikasz-Nagy model (40), this reaction is described by a Goldbeter-Koshland function, which includes the parameter *KA*_*Wee*1*p*_ to regulate the level of Cdc14 required for the phosphorylation of Wee1.

Therefore, we modified these two parameters, *V_si_* and *KA*_*Wee*1*p*_, to ensure that variations of the cycle duration mostly impact on G1 while the length of S phase remains constant. We assumed that the value of the two parameters scales linearly with the duration of the division cycle, *t_cycle_*, between a lower and upper bound, which prevent aberrant behaviour of the cell cycle model in the dynamically changing conditions of the crypt.

V_si_ is scaled according to:

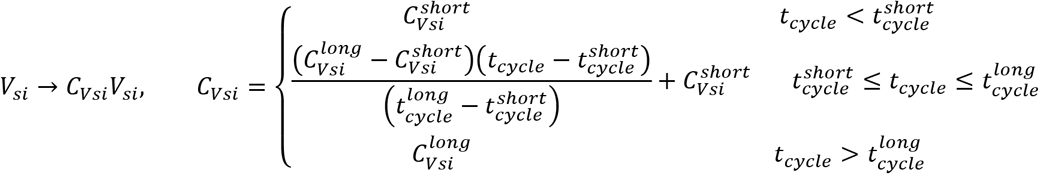

where 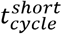 and 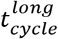 denote the average duration of the cycle of fast cycling progenitors and of the slower cycling stem cells, respectively. 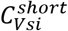 and 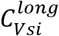 are values calibrated to ensure the correct duration of G1 for the short and long cycle, respectively, and can be found in Supplementary Table 1

Similarly, we scale *KA*_*Wee*1_ using the function:

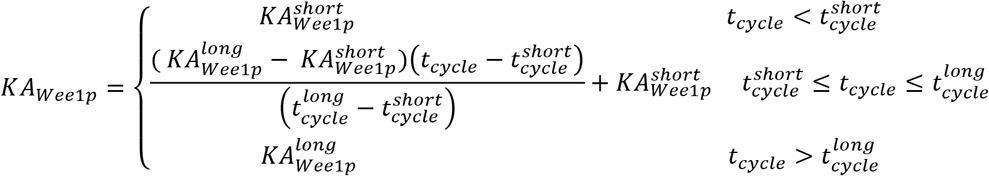

Here 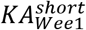 and 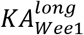 are the values required to maintain constant duration of S-phase in fast and slow cycling cells and can be found in Parameter Table (SX).

A further refinement required to modify the length of the cycle in the Csikasz-Nagy model comprises the mass variable. This variable doubles its value over the course of a cycle and drives the progression of the cell cycle by changing the production rates of the cycle proteins. The changing production rates affect the balance of the proteins and the duration of the cell cycle phases, which start and end at particular mass values determined by the above mentioned two rates and other parameters in the model. After the mass doubles, mitosis occurs and the mass is halved to its initial value, returning the model to the original state. From here the mass begins to grow again, repeating the cell cycle. The mass of a cell effectively tracks the cell’s progress through the cell cycle

In our ABM, *t_cycle_* changes continuously in each cell and modifies *V_si_* and *KA*_*Wee*1*p*_ as described above, which in turn changes the mass values of the start/end of the cell cycle phases. Without further changes in the model, this would cause the cells to not progress through the cell cycle correctly, with unbalanced phases duration and dividing at unwanted mass values, causing erroneous and unrealistic behaviour in the ABM.

This can be solved by normalising the mass in the cell cycle model, chosen such that a cell begins at *mass* = *mass_init_* ≈ 1 and always divides at *mass* = 2.

To do this, we first define a normalised mass variable, assumed to be proportional to the volume of the cell:

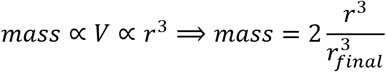

where *r* is the cell radius that takes values between *r_init_* and *r_final_*. When a proliferative cell is created, it is assigned a desired final size, 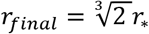, where *r*_*_~*N*(0.35,0.00875) for stem cells and *r*_*_~*N*(0.5,0.0125) for all other cells. In this way, the model captures the smaller apical surface described for columnar stem cells (71), which additionally helps recapitulate the mechanics and cell composition of the niche.

We then introduce a factor *c_mass_* onto the four terms involving the mass variable in the cell cycle model. These terms are the basal production rates of the four cyclins A, B, D and E, called *V_sa_, V_sb_, CycD*_0_ and *V_se_* respectively. *c_mass_* is given by

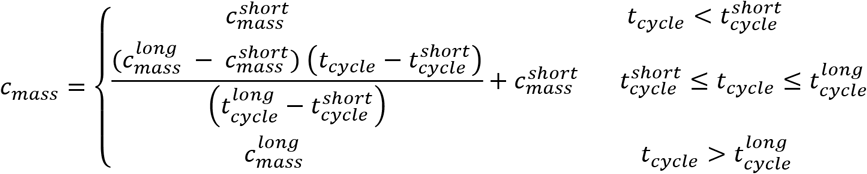

The values 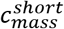 and 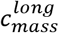 are values found by calibration of the cell cycle model to guarantee the cell always divides at *mass* = 2 for the short and long cycle durations.

Moreover, the cell mass is assumed to grow exponentially. A proliferative cell always reaches a final value of *mass* = 2, corresponding to the radius 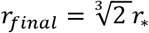, during the cycle time, *t_cycle_*, so that mass must grow as

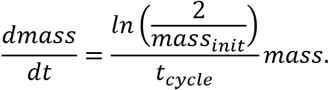

This corresponds to a radial growth rate of

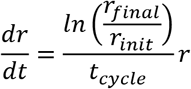

As *t_cycle_* changes dynamically through the cell cycle, the growth rate holds only for the instantaneous conditions the cell is experiencing and changes dynamically through the cell’s lifetime. However, in a healthy crypt extracellular conditions vary slowly, and the value of *t_cycle_* and all derived adjustment factors remain relatively unchanged.

We assumed that cells divide symmetrically. Each daughter cell has a starting radius of 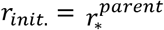 and is assigned with a new randomly generated *r*_*_ value which determines 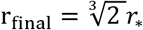. If r_init_. > r_final_, then we set *r_final_* = *r_init_* to prevent values of *mass* > 2. Since cells have a variable maximum size uncorrelated to their birth size, i.e. 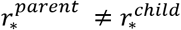, the initial mass value is not necessarily 1. Longer or shorter G1 phases emerge from the model to adjust the cycle duration in cells that begin with *mass* < 1 or *mass* > 1, respectively.

Proliferative daughter cells continue through its own cell cycle and proceed to grow to its own 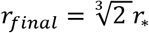. Non-proliferative secretory cells differentiate from stem cells, which are smaller than other cells. To compensate for this, secretory cells grow to reach a radius *r*_*_, generated as *r*_*_~*N*(0.5,0.0125), in a time equal to 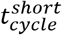. The other type of non-proliferative cells, enterocytes, derive from absorptive progenitors and remain at *r_init_* ≈ 0.5 without increasing size.

These definitions of mass, cell radius and cell growth were chosen to ensure that cells have a consistent radius, and to guarantee that the cell cycle model correctly proceeds through all phases in each cell. Due to the varying cycle duration and extracellular conditions, this control is essential to the correct functioning of the cell cycle and overall behaviour of the ABM.

### 3) Drug perturbations of the Cell Cycle Model: Cdk1 Inhibition

We have used the Csikasz-Nagy cell cycle model to implement drug-induced perturbations of the cell cycle proteins, which are common mechanisms of action of oncotherapeutics, in our ABM. For an arbitrary component of the cell cycle model, *X*, we introduce a term dependent on the drug and *X:*

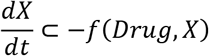

Where ⊂ means “contains the term” and *Drug* represents the cell concentration of the active compound/metabolite which is often described by a pharmacokinetics model. *f*(*Drug, X*), quantifies the effect of the drug on *X*. This function can take several forms such as a mass-action term or a Michaelis-Menten or Hill equation. Multiple terms like this can be added concurrently to the proteins described by the Csikasz-Nagy model.

As an example, we have modelled the effects of a Cdk1 inhibition at the cingle cell level in our ABM. Cdk1 binding is reported to induce nuclear translocation of cyclins A and B require to initiate mitosis (57). Accordingly, we have added a mass-action term onto the rate of change of the CycA/Cdk1/2 and CycB/Cdk1complexes as follows:

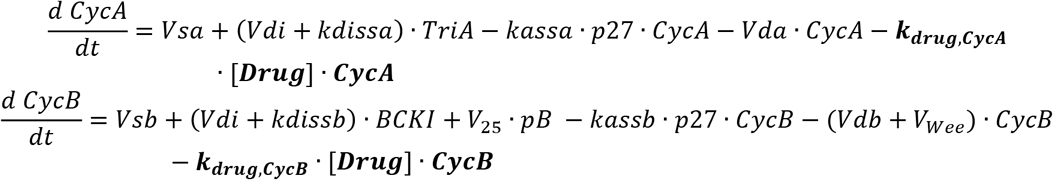

Where *CycA* and *CycB* are used to refer to CycA/Cdk1/2 and CycB/Cdk1 to improve readability of the equation. *k_drug,CycB_ and k_drug,CycA_* are parameters that quantify the drug effect, with values specified in Supplementary Table 1, and [*Drug*] denotes a theoretical drug dynamical concentration.

For the simulation in Figure 4 we assumed that [*Drug*] takes a value equal to 1 for 0 ≤ *time* ≤ 4 *days* and 0 otherwise. Also we considered a smaller the value for *k_drug,CycA_* than for *k_drug,cycB_* to reflect the fact that CycA represents both CycA/Cdk1 and CycA/Cdk2 and only CycA/Cdk1 is inhibited.

These perturbations of the cell cycle proteins can cause incorrect progression throughout the cell cycle, whereupon a cell is permanently arrested. We considered incorrect progression through the cell cycle when the order of the phases was altered, (for example, trying to enter S-phase from M-phase) or phases were skipped. In addition, cells that did not spend sufficient time (30 minutes) in M-phase are killed to recapitulate mitotic catastrophe/death. Note that S-phase is modelled with higher granularity linked to a DNA synthesis model as discussed below.

### 4) DNA and RNA synthesis

Since one of the most common means of targeting the cell cycle is to exploit the effect of DNA-damaging drugs (13), we added the dynamics of DNA replication during S-phase and RNA synthesis during the cell cycle.

Replicating DNA is represented by two variables, *DNA*_1_ and *DNA*_2_, which denote two DNA double helices formed during S-phase. *DNA_i_* is an abstraction of the proportion of undamaged DNA, which takes values from 0, representing total DNA disruption, to 1 for the whole undamaged double helix.

When the cell divides, the daughter cells are given one DNA double helix each (which are both assigned to *DNA*_1_ in the respective cells) to restart the cycle. At the onset of S-phase, the original DNA double helix, *DNA*_2_, unwinds to start the replication of strands and rapidly generates two complete sets of DNA, *DNA*_1_ and *DNA*_2_. This is represented in the model by

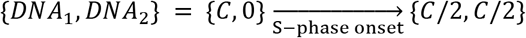

Both *DNA*_1_ and *DNA*_2_ aim at reaching *DNA_i_* = 1. When the cell divides, the daughter cells are given one DNA double helix each (which are both assigned to *DNA*_1_ in the respective doughtier cell) to restart the cycle. Outside S-phase, DNA synthesis takes place solely for repair at a slower rate.

Hence, in healthy cells, these variables obey the following equations and algorithm:

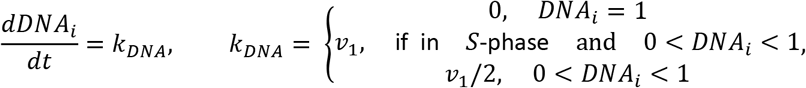

The DNA replication rate, *v_1_*, is sufficiently fast to ensure *DNA_i_* reaches 1 during S phase in healthy cells. Outside of S-phase, we assumed a 2-fold slower rate for DNA repair when the cell is not actively replicating its DNA. Values are specified in Supplementary Table 1.

RNA levels are represented by a single *RNA* variable. Similarly, this variable is an abstraction of the proportion of undamaged RNA in the cell, with *RNA* = 1 in a healthy cell and *RNA* = 0 for total RNA disruption. RNA synthesis is assumed to be governed by a simple linear-growth differential equation until its maximum value, *RNA* = 1, and remains at this value unless damage is induced as follows,

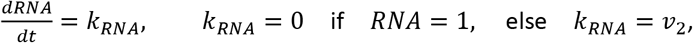

with parameter values specified in Supplementary Table 1.

Along with these equations for DNA and RNA levels, we added DNA and RNA-damage checkpoints to modulate the response of the Csikasz-Nagy cell cycle model to perturbations. We considered both the G1/S and the G2/M checkpoints (44), with cells checking their DNA and RNA levels as they progress from G1 to S-phase, and from G2 to M-phase. If the DNA and/or RNA levels are below the threshold values (see Supplementary Table 1), the cell undergoes apoptosis. Checkpoint failures can occur upon drug-induced DNA or RNA damage, as explained below.

### 5) Drug perturbations of RNA and DNA synthesis: 5-FU induced RNA and DNA damage

Similar to the cell cycle model, drug effects are represented by adding a negative term to these differential equations:

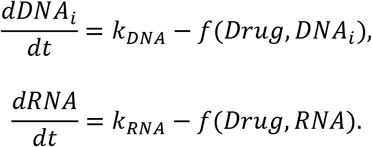

Where *f*(*drug, X*) could be a mass action, Hill equation or Michaelis-Mentin term quantifying the drug induced RNA or DNA damage.

DNA damage induces increased p21 expression in cells, which prevents progression through the cell cycle and can lead to cell cycle arrest or apoptosis (72). To replicate this, we further modified the p21/p27 term in the Csikasz-Nagy model to respond to the DNA levels of the cell. Recall that *V_si_* was the production rate of p21/p27 in the model, and we multiplied this by *C_Vsi_* to moderate the production of p21/p27 (see details in ‘Cell Cycle Proteins and Cell Size/Growth’ above).

Recall that *V_si_* → *C_Vsi_ V_si_*, to replicate DNA-damaged induced production of p21, we replace *C_Vsi_* with a bounded function dependent on the cells’ DNA levels

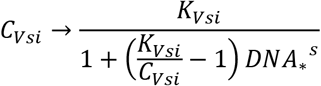

*DNA*_*_ = *DNA*_1_ in G1, and *DNA*_*_ = *min*(*DNA*_1_ + *DNA*_2_, 1) in all other phases and *s* is a scaling coefficient. In homeostasis, with *DNA*_1_ + *DNA*_2_ ≥ 1, this function is equal to *C_Vsi_* and the cell cycle model proceeds as before. With severe DNA damage, *DNA*_*_ << 1, the function is approximately equal to *K_Vsi_*, always > *C_Vsi_*, that represents the maximum fold increase of the production rate of p21, i.e. *V_si_* → *K_Vsi_ V_si_*. Parameter values can be found in Supplementary Table 1. When DNA levels are reduced by drug-induced injury, this new function increases the production rate of p21/p27 which slows down the production of cyclins and the progression of the cell cycle, recapitulating a reversable cell cycle arrest for low-to-moderate DNA damage (73).

Cell growth is dependent on the correct translation of mRNA into proteins. We hypothesised that RNA damage reduces a cell’s capability of biosynthesis and leads to slower cellular growth (74). This is modelled by adding an RNA-dependent factor to the growth rate of cells:

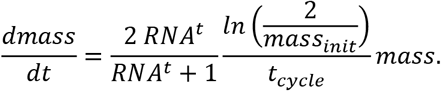

Where *RNA* takes as defined above values between 0 and 1 and *t* is a scaling coefficient. Parameter values can be found in the Supplementary Table 1. By linking RNA integrity to cellular growth, we allow RNA damage to induce a form of cell cycle arrest, as previously reported (75, 76).

The result of these responses to DNA and RNA damage, in combination with the cell cycle checkpoints, allows the cells in our model to exhibit a progression of responses to increasingly severe DNA and RNA damage. Cells with low damaged DNA and/or RNA levels grow and proliferate slowly, due to impediment of their cell cycle and/or cellular growth. With moderate DNA and RNA damage, a cell enters an impermanent, reversible cell cycle arrest (characterised by a near zero growth rate and p21-induced halt of the cell cycle). Upon interruption of the drug-induced insult, these cells will re-enter the cell cycle. In case of severe DNA and/or RNA damage, a cell will undergo DNA/RNA damage-induced apoptosis caused by failing a cell cycle checkpoint. Additionally, drug-induced perturbations may result in incorrect progression through the cell cycle, which causes the cell to enter a permanent arrested state or die as described above. Note that though RNA damage is known to cause cell cycle arrest and apoptosis (76), the mechanisms are poorly known, so we made the conservative decision to check the level of RNA damage at the same checkpoints as DNA damage.

As an example we modelled 5-FU induced RNA and DNA damage in the intestinal epithelium. We considered the two main downstream metabolites of 5-FU, FdUTP and FUTP, causing DNA and RNA damage, respectively (62). To do this, we implemented in the ABM a previously published model that describes 5-FU distribution post-dosing in mouse and a reduced version of the 5-FU metabolic pathway(65).

Furthermore, we implemented the effect of FdUTP and FUTP on DNA and RNA synthesis, respectively, on each cell of our ABM using a Hill function as follows,

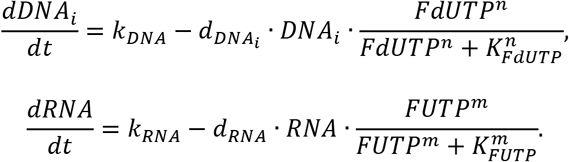

Parameter values can be found in Supplementary Table 1.

The impact of these metabolites on DNA and RNA of each cell of the epithelium resulted in the arrest of the majority of proliferative cells, with a small proportion undergoing apoptosis after failing the G1/S or G2/M checkpoint.

### 6) Mechanical Cell Interactions and Contact Inhibition

Intestinal stem cells and early progenitor cells compete for limited niche space and, therefore, the ability to retain or regain stemness. Cell proliferation creates a constant battle for space, inducing forces that drive cell migration away from the hard boundary of the stem cell niche towards the top of the crypt and onto the villus.

We assumed intercellular physical forces based on Hertzian contact mechanics with adhesive and frictional forces, similar to those in published reports (77) (17). For the sake of simplicity and differently from previous approaches, we did not include the extra repulsive force opposing the reduction in cell volume caused by cell overlapping and did not consider radial expansion of cells to compensate for the loss of volume in compressed cells.

In our model, cells experience repulsive, adhesive, and frictional forces. Forces results in movement according to Stoke’s flow, where viscous forces dominate inertial forces, and cell velocity is directly proportional to the resultant forces on the cell. For very shallow overlapping distances (^ 10 *%* of the cells radius), the adhesive force holds the cells together and replicate continuity of a biological tissue, but for greater overlap distances, repulsive forces dominate. Frictional forces help create collective movement by counteracting cell migration in the opposite direction to the general flow of cells.

All distances are expressed in arbitrary cell units defined such that the diameter of an average cell is equal to 1. Forces are then measured in the resulting units.

Below we describe these forces and porceses in detail.

#### 6.1. Contact Repulsion

Cells are assumed to be elastic spheres with inter-cellular forces derived from Hertzian contact mechanics. The magnitude of the repellent force, 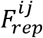. between cell *i* (with position vector **x***_i_*, radius *R_i_*, Young’s modulus *E_i_* and Poisson ratio *ν_i_*) and cell *j* (with position vector **x***_i_*, radius *R_j_*, Young’s modulus *E_j_* and Poisson ratio *ν_i_*) is described as follows

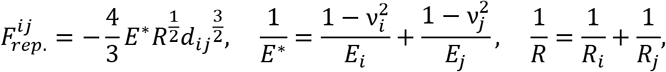

where *d_ij_* = *R_i_* + *R_j_*n |**x***_ij_*| is the overlapping distance between cells measured on the line joining the cell centres, with **x***_ij_* = **x***_j_* – **x***_i_* the displacement vector joining the two cell centres. This repulsive force acts on both cells in opposing directions, pushing them away along the unit vector joining the two cells 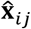:

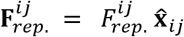

The reported value for the Young’s modulus of Paneth cell is relatively large (35) and results in a relatively large force acting on neighbouring stem cells which helps to confine them in the niche. In addition, the previously published values of the Poisson ratio indicate that cells are marginally compressible(78).

#### 6.2. Adhesive Force

All cells in contact experience adhesive forces proportional to the area of contact and the cells inherent adhesiveness, parameterised by ε. The magnitude of adhesive force between cell *i* and *j* is quantified as follows

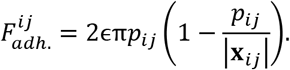

where |**x***_ij_*| is the distance between cell centres and

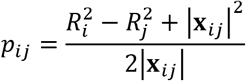

This force is again directed along 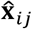, pulling the cells together: 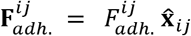 and its magnitude is derived by assuming the associated energy, 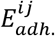, is proportional to the area of contact between cells *i* and *j*, 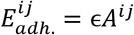, where 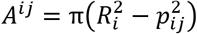, and differentiating with respect to the distance between the cells.

Two cells in isolation will be at rest when the repulsive and adhesive forces are equal, however in our simulations, this rarely happens due to the constant proliferation and growth of surrounding cells. In vivo crypts have a highly compressed niche with tightly packed stem cells wedged between Paneth cells. In our model, the repulsive force is parameterised entirely by observed quantities (the Young’s modulus and Poisson ratio), leaving *ϵ* in the adhesive force as a free parameter. The value of *ϵ* determines intercell separation at rest. This value was chosen to allow overlapping of Paneth cells at rest of 0.15 distance units, which corresponds to 15% of the diameter of an average Paneth cell. This results in *ϵ* = 0.216 for Paneth-Paneth adhesion. Qualitatively, all other cells are less tightly packed, so all other adhesive forces (including Paneth cells with any other cell type) are assumed to be 10-fold weaker with *ϵ* = 0.0216, which produces an overlap of approximately 0.075 cell units. These assumptions facilitate the recapitulation of the tighter packed cells in the niche resulting in increased mechanical pressure (defined in the following sections) which induces proliferation contact inhibition mechanisms.

#### 6.3. Frictional force

Cells moving with a relative velocity while in contact experience a frictional force. The force acting upon cell *i* due to friction with cell j is quantified as follows:

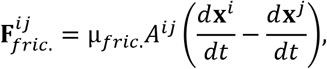

where *A^ij^* is the area of contact between cells *i* and *j* defined above, and *μ_fric_*. is a numerical constant calibrated to enforce orderly cell dynamics. This force is quite small but helps collective motion of cells by opposing cell migration against the common direction.

#### 6.4. Cell migration

Under a force, cells move according to Stoke’s flow, where viscous forces are assumed to dominate over inertial effects:

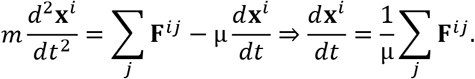

Therefore, the position vector of the *i*-th cell, **x***^i^*, is updated according to

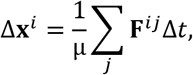

where 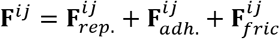 is the resultant of all forces on cell *i* due to cell *j*.

The parameter μ links the forces to cellular motion. The value of this parameter is estimated to recapitulate the transfer velocity in the crypt-villus junction measured in in vivo experiments to be approximately 1 cell position per hour in mice (79). However, cell motion response to these forces may vary for different cells types. It has been reported that that Paneth cells persist in the stem cell niche at the crypt base for relatively long periods of up to 57 days in mice (80, 81) and exhibit elevated β_4_-integrin expression anchoring them to the mesenchyme (82). Additionally, Paneth cells are larger and stiffer than the comparatively malleable stem cells which suggest that they require greater forces to be displaced. In our model, we used μ to replicate this behaviour and recreate drag effects of the basal membrane/mesenchyme. We implemented a value of μ for Paneth cells 10000-fold greater than for other cells, effectively making Paneth cells difficult to move by other cells but allowing them to slowly move one another to form an orderly niche over longer timescales.

#### 6.5. Internal Pressure and Contact Inhibition

The forces described above are used to calculate the internal pressure experienced by cells, which varies according to the cell intrinsic properties and local environment, i.e. a stem cell in the crowded niche, has higher internal pressure. Cell pressure is used to recapitulate contact inhibition by modulating the duration of the division cycle which increases when cells are densely squeezed together and decreases if cell density falls to enable, for instance, fast recovery from injury.

A cell feels internal stress from the surrounding cells, and this is used to simulate contact inhibition. To do this we use the concept of virial stress outlined in (83). The stress tensor for cell *i, σ_i_*, is defined as follows:

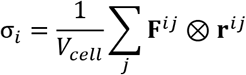

where **r***^ij^* is the vector from the centre of the cell *i* to the plane of contact with cell *j*, always assumed to be on the surface of cell *i*, and ® is the tensor/outer product combining two vectors into a ‘matrix’. Using this stress tensor, we extract the pressure in the conventional manner:

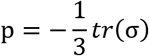

As all our forces are normal to the plane of contact, this reduces to

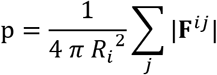

This provides a rough, first-order approximation to the pressure experienced at the centre of the cell that is straightforward to compute and essential to implement contact inhibition in proliferative cells. Note that we do not consider the hydrostatic pressure induced by cell compression.

On the other hand, physical compression has been reported to lead to YAP inactivation, retarding growth and morphogenesis in the GI epithelium (32–34). We used our estimate of pressure to implement this contact proliferation inhibition mechanism responding to the mechanical environment and described the increase of the cell cycle duration, *t_cycle_*, as pressure, *p*, increases using a scaled logistic function as follows

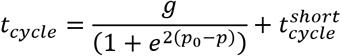

Here *p*_0_ is the average pressure experienced by cells in the niche; 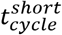 is the average division time of absorptive progenitors and 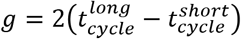 where 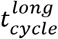 denotes the longer division time of a stem cell in average niche conditions.

This function captures the variation of the duration of the division cycle from a minimum to a maximum value in cells highly compressed which leads to longer division times in the tightly constrained stem cell niche of the crypt, while the cycle is shorter in the less compressed transit amplifying zone, in agreement with experimental reports (4) (84) (85).

### 7) Biochemical Signalling

Next, we detail how the cells interact with one another, communicating the local composition of the crypt to maintain homeostasis. This is done through simulated biochemical signalling

To achieve stable crypt cell composition and structure, we have implemented five signalling mechanisms including Wnt, Notch and BMP pathways which have been demonstrated to be essential for morphogenesis and homeostasis of the intestinal crypt (20) (21) (22) (23) (5). We have modelled contact proliferation inhibition mediated by the YAP-Hippo signalling pathway responding to mechanical forces (32–34) as described above and following experimental evidence (7) (6) (25), implemented a RNF43/ZNRF3-like mediated feedback mechanism between Paneth and stem cells.

These minimal signalling mechanisms were chosen because a full understanding of the protein interaction networks is still a topic of active research. However, even with our conservative assumptions, we implicitly introduce crosstalk between the different signalling pathways. For example, the nature of cell fate decisions leads to interaction between Wnt and Notch levels, and changes in the duration of the cell cycle caused by contact inhibition regulates the ability of a cell to accumulate signalling molecules.

#### 7.1. Wnt signalling

The Wnt pathway is the primary pathway associated with stem cell maintenance and differentiation in the intestinal crypt as well as in many other tissues (26) (20) (86). Two sources of Wnt signals have been described in the mouse crypt: Paneth cells (27) and speficific mesenchymal cells surrounding the stem cell niche at the crypt base (28).

We did not consider the dynamics of the canonical Wnt signalling molecular cascade but directly implemented downstream cellular responses to Wnt levels. We modelled Wnt signalling as a short-range (1 cell diameter) field around Paneth cells and Wnt-emitting mesenchymal cells at the bottom of the crypt and considered Wnt signals tethered to receptive cells as previously reported (25, 29). This is described by the following equation:

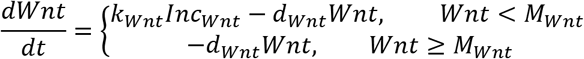

where, pictorially, we have

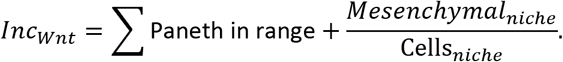

The variable ‘*Wnt*’ is an abstraction of the total number of Wnt ligands tethered to the surface of the cell; *Mesenchymal_niche_* represents the number of Wnt emitting mesenchymal cells surrounding the niche, which we assume is equal to the total number of epithelial cells in the niche in homeostatic conditions, *Cells_niche_*. Additional Wnt production by Paneth cells is required to support the homeostatic number of stem cells in homeostasis. In the presented modelling scenarios, we assumed constant exogeneous Wnt source, i.e. constant *Mesenchymal_niche_*, shared by all cells in the niche and enhancing niche recovery after damage. For instance, with lower number of cells in the niche, the survival cells will receive stronger mesenchymal Wnt signalling that enhances proliferation and recovery after perturbations. We assumed that surface tethered signals are equally distributed between daughter cells upon cell division (5, 25), so that cells eventually lose Wnt signals and their capacity to proliferate if not within the range of a Wnt source. These assumptions are partly supported by observed in vivo and in vitro behaviour, where the mesenchymal and Paneth cell derived Wnt sources are mutually redundant (87).

The variable *M_Wnt_* describes the maximum number of Wnt signals a cell can have tethered and is given the arbitrary value of 128 in all cells. We chose values to be powers of 2, i.e. 64 and 128, to facilitate dividing Wnt signals in half upon cellular division.

#### 7.2. ZNRF3/RNF43 Signalling

In our model Paneth cells enhance their own production by generating high Wnt local environments (88). In addition, due to their high Young’s modulus, Paneth cells create a region of high intercellular forces on neighbouring cells which leads to prolonged division times with greater opportunity for Wnt accumulation. This in turn expands the niche region with high Wnt and high cell pressure, promoting further differentiation into stem and Paneth cells. Therefore, without a negative feedback mechanism in our model, these features would result the expansion of the niche with stem and Paneth cells occupying the entire crypt. Additionally, two recent studies have demonstrated the existence of a negative feedback loop mediated by RNF43 and ZNRF3 ligands produced by stem cells (6, 7). These studies proposed that RNF43 and ZNRF3 inhibit Wnt signalling by promoting the turnover of Wnt receptors such as Frizzled and LRP5 (89), and showed that simultaneous deletion of these two receptors results in the formation of adenomas comprising mostly stem and Paneth cells (7).

We assumed that RNF43/ZNRF3 (henceforth called ZNRF3 for simplicity) is a diffusing, decaying signal secreted by stem cells. Without explicit knowledge of the chemical and physical properties of ZNRF3 signalling, this process is assumed to immediately reach steady state at the timescale of cellular decisions. Therefore, the ZNRF3 signal strength, *ZNRF3(r)*, received by a cell at position *r* from a stem cell located at position *R*. is described by the diffusion equation as follows (90)

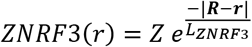

where *Z* represents the maximum signal strength immediately around the emitting cell, for simplicity we assumed *Z* =1. *L*_*ZNRF*3_ determines the spatial scale of diffusion, which we assumed is equal to the length of a cell in order to maintain high signalling levels primarily in the niche.

The total ZNRF3 signalling received by a cell at position *r* is calculated, therefore, as the sum of the signal received from all stem cells:

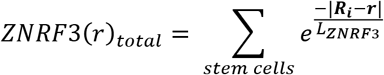

where *R_t_* the position of the *i*-th stem cell.

The strength of ZNRF3 signalling received by a cell is proportional to the number of stem cells in the immediate vicinity of the cell: in typical, homeostatic conditions, *ZNRF3*(*r*)*_total_* is high in the niche, falling off exponentially as a cell moves towards the villus.

The ZNRF3 signalling level detected by a cell, located at position ***r***, regulates the decay rate of its surface-tethered Wnt molecules, *d_Wnt_* as follows:

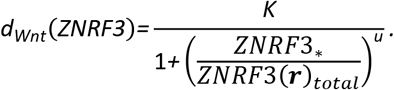

where *u* is a scaling coefficient, and *K* and *ZNRF3*_*_ are constants calibrated to maintain the size of niche at its homeostatic level. In particular, *ZNRF3*_*_ is determined by the homeostatic number of stem cells in the niche, while *K* was calibrated to produce a Wnt decay rate high enough to prevent Wnt values ≥ 64 in cells located at the edge of the niche when the number of stem cells is excessive such that *ZNRF3*(***r***)*_total_* ≥ *ZNRF3*_*_. These considerations prevent the expansion of the niche by preventing cells from differentiating into the Paneth or stem cell fate (which requires Wnt ≥ 64) when a cell is outside the niche.

With this implementation of ZNRF3-mediated negative feedback, the Wnt decay rate within the niche is high but is compensated by the abundant Wnt supply from mesenchymal and Paneth sources, while the Wnt decay rate rapidly drops to zero outside the niche. This means that the degradation of Wnt outside the niche has little impact on a healthy crypt and the Wnt gradient in our model is mainly generated by the halving of the surface bound Wnt signals between daughter cells upon division. Growth and proliferation derived forces drive migration of cells towards the villus while the amount of tethered Wnt decreases after each division. This recreates the observed (25) decreasing gradient of Wnt signals moving up the crypt (Figure 1), with the highest values in the niche, intermediary values in the transit-amplifying zone and low levels in the upper crypt region of differentiated enterocytes.

The stem cell-mediated negative feedback loop regulating Wnt signalling, together with the differentiation rules described below, ensure the maintenance of the niche size and crypt composition in homeostasis. In addition, it also facilitates crypt recovery as stem cells in low numbers are able to reach greater surface-tethered Wnt levels to pass to their offspring which, in turn, can more readily acquire the required amount of Wnt to become stem cells.

#### 7.3. Notch signalling

Active Notch signalling requires direct membrane contact between two cells, one expressing Notch ligands and the other Notch receptors (91) (22) (92) (5). In the intestinal epithelium, Notch ligands present in secretory cells bond to transmembrane notch receptors of stem cells to induce a transcriptional cascade which blocks differentiation of stem cells into the secretory lineage in a process known as lateral inhibition and leads to checkerboard/on-off pattern of Paneth and stem cells in the niche (21) (93). With these considerations, Notch signalling is implemented according to the following equation:

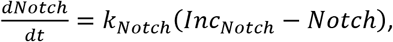

where *Inc_Notch_* is the amount of incoming notch ligands to the cell which we assumed is equal to the number of ligands expressing cells in contact with the cell. At steady-state, a cell’s Notch value corresponds to the number of incoming Notch ligands the cell is receiving: for example, a stem cell receiving Notch from one single neighbouring cell, reaches equilibrium with *Notch* = 1. The factor *k_Not_ch* denotes the rate of Notch accumulation and it has a relatively high value to ensure that the equilibrium is reached before the fate-commitment point at the end of G1. As described in the cell cycle section, the duration of G1 changes with the length of the overall division cycle: shorter cycles have a shorter *G*_1_ phase, shortening the time the cell has to receive Notch signals before deciding whether to differentiate or divide. Additionally, *k_Notch_* is also the decay rate of the cell Notch signalling and this relatively fast rate means that Notch must be constantly supplied for a stem cell to maintain stemness.

A reduction in cell density (e.g. by ablation of cells) can introduce gaps in the simulated epithelial tissue. In real tissues, these gaps would be covered by expansion-flattening of surviving cells to restore epithelial integrity and contact to neighbouring cells. These new contacts would allow cells to exchange Notch ligands. In our model, we do not explicitly consider the expansion of cells to fill gaps in the epithelium, however, we simulate this effect by allowing a cell to pass Notch signals to receiving cells within a larger range (1 cell diameter) following a drop in local density. This allows our model to recreate the correct recovery response following ablation of cells.

#### 7.4. BMP signalling

The Wnt gradient in the crypt is opposed by a gradient of bone morphogenic protein (BMP) generated by mesenchymal telocytes, which are especially abundant at the villus base and provide a BMP reservoir, and by the recently identified trophocytes located just below crypts and secreting the BMP antagonist Gremlin1 (37). BMP signals inhibit cell proliferation and promote terminal differentiation (36). Large levels of BMP at the crypt-villus junction prevent proliferative cells from reaching the villus (94). BMP signalling has been reported to be modulated by matured epithelial cells on the villus via hedgehog signalling (10, 11) such that a decrease of villus cells decreases BMP signalling in the crypt, which enhances proliferation and expedites villus regeneration.

We propose a simple model that assumes that enterocytes, *E*, secrete diffusing signals, which could be interpreted as Indian Hedgehog, to regulate BMP secretion by mesenchymal cells. The explicit pathways and associated timescales involved in BMP signalling are unknown, therefore, similar to our implementation of ZNRF3 signalling, this process is assumed to instantaneously reach steady state at the timescale of cellular decisions. As before, we assume that BMP is a diffusing, decaying signal in steady state (90) described by

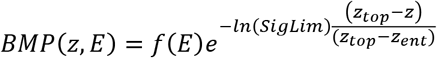

where *z* is the position coordinate corresponding to the crypt-villus longitudinal axis; in our model *z ≤* 0 for cells located in the stem cell niche while z > 0 for crypt cells outside the niche; *z_top_* is the maximum z value at the top of the crypt; *SigLim* is the exponential transformation of the diffusion coefficient. To facilitate the use of the model for different species, the *z* coordinate is standardized using *z_ent_*, which is the crypt axis position at which the number of mature enterocytes becomes greater than the number of absorptive progenitors. As mentioned above, mesenchymal cells surrounding the niche secrete BMP antagonists (37) and we assumed that BMP signalling is effectively blocked in the niche such that *BMP* = 0, which is approximately true for the above formula. *f*(*E*) describes the relationship between the number of enterocytes and maximum BMP signal intensity using an increasing Hill function:

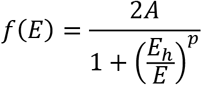

where *E_h_* is the homeostatic number of enterocytes determined by in-vivo experiments, *p* is the Hill interaction coefficient and *A* denotes the level of BMP signals at position *z_top_*. In our model, absorptive progenitors differentiate into enterocytes when *BMP* > *Wnt*, representing that the anti-proliferative BMP signalling received by the cell is sufficient to overcome the proliferative effect of Wnt (23). We achieved an optimal crypt cell composition with values of *A* and *SigLim* that allow progenitors cells to divide in a healthy crypt at least 3 times before differentiating. Differentiation occurs when the cell Wnt content reaches values below *SigLim* = *BMP*(*z_ent_*) when migrating upwards.

In addition, these equations describe a frequently reported feedback response to villus injury consisting of enhanced proliferation within hypertrophic crypts (10, 38, 39). In our model, when the number of enterocytes on the villus falls below the homeostatic level, the production of BMP signals decreases and makes it possible for absorptive progenitors to divide more times and reach higher positions in the crypt before becoming terminally differentiated. Concurrently, the height of the crypt must increase to provide sufficient space for the extra proliferative cells. We modelled the enlargement of the crypt height, *h*, responding to villus injury using a decreasing Hill function as follows:

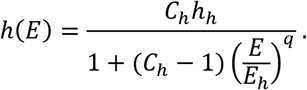

where *h_h_* is the observed crypt height in homeostasis, *C_h_* is the maximum fold increase of the height of the crypt and *q* the Hill interaction coefficient. We do not consider cases in which the number of enterocytes on the villus increases above homeostatic levels, such that if *E* > *E_h_* then *h*(*E*) = *h_h_*

### 8) Cell fate: proliferation, differentiation, arrest, apoptosis

In the sections above, we have outlined the dynamics of signalling pathways, cell cycle proteins and mechanical forces. These processes interact with each other to maintain epithelial homeostasis by precisely tuning cell proliferation, differentiation and migration within the crypt geometry.

An overall picture integrating the rules governing cell fate decision is described in Fig 1. Wnt levels ≥ 64 arbitrary units (au) are required for stemness maintenance. For a stem cell, lateral inhibition is repressed when Notch < 3 au, equivalent to less than 3 secretory cells in the local neighbourhood. If Notch is repressed (<3 au) and Wnt >64 au, stem cells differentiate into Paneth cells. Paneth cells generate Wnt signals which enhance the production of stem cells and of Paneth cells themselves. Niche expansion is modulated by the RNF43/ZNRF3 mediated negative feedback mechanism (25) (6) (7) that makes Wnt>64 unobtainable after reaching the homeostatic number of stem cells. Furthermore, the duration of the division cycle is dependent on local forces experienced by the cell. Cells under high mechanical pressure (in the niche) are subjected to YAP-Hippo regulated contact inhibition and with longer cycles accumulate more Wnt and Notch signals. On the other hand, cells located outside the niche exhibit shorter cycles and cannot effectively accumulate enough Wnt signals to become stem or Paneth cells.

Stem cells with decreased levels of Wnt signalling (<64), usually located outside the niche, differentiate into absorptive proliferating progenitors if Notch signalling is active or into secretory progenitors if Notch signals < 2 au. This lower Notch threshold value is required to maintain the correct balance of absorptive and secretory cells outside the niche, in the absence of large numbers of Notch secreting Paneth cells. All cells migrate towards the crypt mouth driven by proliferation forces. During this migration the Wnt content in absorptive progenitors is halved in each division and, away from Wnt sources, progressively decreases while BMP levels progressively increases towards the villus. Eventually, progenitors reach reduced Wnt content and increased BMP levels required for differentiation into enterocytes.

All fate decisions are assumed to be made at the restriction point which in our model is located at the end of G1 (95). At the restriction point, cells assess their internal Wnt and Notch levels and if these values fulfil the criteria to differentiate they enter a quiescent state or G0, otherwise they proceed to S-phase and become irreversibly committed to complete the cell cycle of variable duration depending on local forces. This quiescent state lasts for 4h for all differentiating cells, except for absorptive progenitors, which differentiate straightaway into enterocytes. In accordance with (96), a secretory progenitor requires an additional 4h to fully mature into a goblet or enteroendocrine cell.

Therefore, the so-called +4 stem cells (97) (98) (5) emerge naturally in the model as stem cells migrate outside the niche and pause the cycle to give rise to non-proliferative secretory progenitors, which have been identified with quiescent +4 stem cells (53) (99). Features and behaviours of these cells could be expanded if of interest for the model application.

Cell fate decisions are reversible; a stem cell that leaves the niche and differentiates into a progenitor cell can relatively quickly become a stem cell again if regaining enough Wnt signals by being pushed back into the niche. This plasticity extends to all cells: all progenitors and fully differentiated cells can revert to stem cells when exposed to sufficient levels of Wnt and Notch signals, replicating injury recovery mechanisms observed in the crypt (48, 51). We have assumed that all cells, except Paneth cells, need to acquire and maintain high levels of Wnt signals (>64) over 4 hours to complete the process. Dedifferentiating cells shrink to their new smaller size during the process if required.

Notch signalling mediates the process of Paneth cell de-differentiation into stem cells to regenerate the niche as previously reported (30, 31). Paneth cells not supplying Notch ligands for 12h to recipient cells dedifferentiate into stem cells in a process that takes 36h to complete in agreement with published findings (31).

Additionally, Paneth cells in low Wnt conditions (for example, a Paneth cell that is forced out of the niche) for 48h will also dedifferentiate into a stem cell, which with low Wnt content rapidly becomes a secretory or absorptive progenitor.

Additionally, injured proliferative cells can experience cell cycle arrest and apoptosis, induced by drug injury or by natural senescence. In arrested and apoptotic proliferative cells, the production rates of the cyclins (*V_sa_, V_sb_, CycD*_0_ and *V_se_*) are set to 0 to interrupt the cell cycle. We assumed that cells remain arrested until they are shed from the villus tip or reach the end of their lifespan and become apoptotic. Apoptotic cells shrink and die with a negative linear rate of

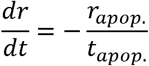

where *r_apop_* is the radius at the onset of apoptosis and *t_apop_* is the time for the completion of apoptosis.

### 9) Model implementation and parametrization

The model is implemented using the Julia programming language. The mechanical forces, cellular motion and biochemical signalling are simulated with a fixed timestep of *dt* = 0.0001 days, while the proteins of the cell cycle model are simulated with a timestep of 0.00001 days. Parameter values and means used for their identification are detailed in Supplementary Table 1.

